# Light-dependent switching of circling handedness in microswimmer navigation

**DOI:** 10.1101/2025.08.08.669292

**Authors:** Zhao Wang, Samuel A. Bentley, Jiawei Li, Kirsty Y. Wan, Alan C. H. Tsang

**Affiliations:** Department of Mechanical Engineering, The University of Hong Kong, Pokfulam Road, Hong Kong, China; Living Systems Institute & Department of Mathematics and Statistics, University of Exeter, Exeter EX4 4QD, United Kingdom

## Abstract

Many swimming microorganisms navigate their environment by modulating the curvature of their swimming trajectories in response to external cues. Here, we show that the biflagellate alga *Chlamy-domonas reinhardtii*swims in circles and actively switches its trajectory handedness in response to orthogonal illumination: the cell swims counterclockwise at low light intensities yet clockwise at high light intensities. This handedness switching arises from light-dependent modulation of flagellar beating, including changes in beat extension, phase, and—crucially—beat plane orientation. Using high-speed imaging and hydrodynamic modeling, we reveal that this beat plane reorientation is critical for *Chlamydomonas* to swim orthogonally to light as well as to dynamically modulate its trajectory curvature, enabling transitions between global exploration and localized searching in spatially structured light fields. Our results establish beat plane reorientation as a novel mechanism for curvature control in microswimmer navigation.

---

Navigating complex and dynamic environments is essential for the survival of biological microswimmers [1–3]. Different species have evolved distinct navigation strategies [4–9], such as the canonical “run-and-tumble” (RnT) behavior in *Escherichia coli* [10], where cells alternate between straight swimming and abrupt reorientation. Another example is the continuous curvature modulation observed in sea urchin sperm [11], which allows smooth steering toward chemical cues. Mammalian sperm can even switch between these two strategies depending on the rheological properties of the medium [12]. Despite these diverse behaviors, the biophysical principles by which sensorimotor coupling enables switching between versatile navigation strategies remain poorly understood.

Many microswimmers actively couple helical swimming with self-rolling to scan the environment in three dimensions [13–18]. They typically exhibit RnT in dim environments, with the run phase being relatively straight and symmetric in the absence of external stimuli. However, environmental cues such as chemical gradients, physical boundaries, or light stimuli can break this symmetry and alter trajectory curvature. For example, proximity to surfaces can induce clockwise (CW) or counterclockwise (CCW) circling due to hydrodynamic effects [19–21], while chemical gradients can trigger asymmetric flagellar beating and CCW swimming in sea urchin sperm [11]. In response to strong physical confinement, *Chlamydomonas* cells exhibit non-equilibrium fluxes in their motion trajectories with a light-dependent handedness [22, 23]. However, the mechanistic basis of how microswimmers actively modulate trajectory curvature or handedness in response to environmental stimuli, and how this guides navigation, is still not well understood.

In this Letter, we report a light-induced circular swimming behavior in *Chlamydomonas reinhardtii*, where the cell modulates trajectory curvature and handedness in response to light intensity. When *Chlamydomonas* cells are exposed to light incident from above [light vector in *−z* direction, Fig. 1(a)], they transition from straight helical swimming in darkness (~ 0 lx) to CCW circular trajectories at low light (~ 150 lx), and subsequently transitions to CW trajectories at high light (~ 15, 000 lx). This handedness switching occurs in bulk far from boundaries in deep chambers (*≥* 100 *µ* m depth), confirming it is not due to hydrodynamic interactions with surfaces. We reveal that this behavioral switching arises from light-dependent modulation of flagella beat pattern and beat plane reorientation. We conclude that coordination of these light-intensity dependent behaviors enables gradient ascent, local and global exploration in spatially structured light fields, providing a new perspective on the role of trajectory curvature modulation in microswimmer navigation.

**FIG. 1.**
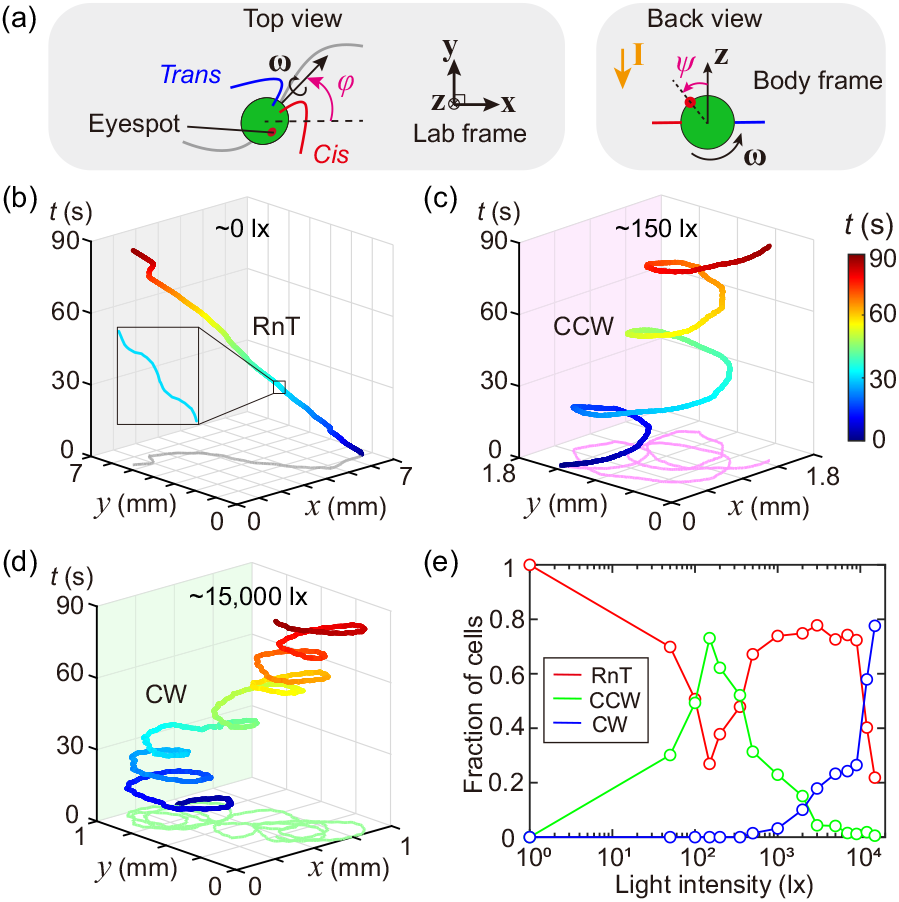
*Chlamydomonas* switches trajectory handedness in response to light intensity. (a) Schematic of *Chlamydomonas* swimming in a helical trajectory with left-handed self-rolling (angular velocity ***ω***). (b) In darkness (~ 0 lx), cells display a RnT trajectory (Movies S1). (c) At low light (~ 150 lx), cells swim in CCW circles (Movie S2). (d) At high light (~ 15,000 lx), cells switch to CW circling (Movie S3). (e) Fraction of behavioral states across light intensities in *Chlamydomonas*. More than 2,000 cells were tracked in total.

## Handedness switching

*Chlamydomonas reinhardtii* is a unicellular green alga that swims using two anterior flagella and senses light with a single photoreceptor organelle called the eyespot. The eyespot sits on the equator of the cell body, offset by 45° from the flagellar beat plane, closer to the *cis-* flagellum [24] [Fig. 1(a)]. We characterize cell swimming dynamics by defining *φ* (cell orientation in the lab frame) and *ψ* (eyespot position in the body frame). Unlike typical phototaxis, where the cell swims toward or away from light [25–30], we focus here on photoresponses to orthogonally directed light, with the swimming plane perpendicular to the stimulus direction. By tracking free-swimming cells in fluid chambers (~ 2 cm *×* 2 cm *×* 100 *µ* m) over extended periods (~ 5 minutes), we reconstructed and analyzed their trajectories in the lab frame (see Supplementary Section 2.2).

In darkness (~ 0 lx), *Chlamydomonas* follows nearly straight helical trajectories interrupted by abrupt, large reorientations, resembling eukaryotic RnT behavior [31] [Fig. 1(b)]. Upon light stimulation, the cell’s trajectory becomes circular, with the handedness depending on light intensity: at low intensity (~ 150 lx), the cell swims along CCW circles; at high intensity (~ 15,000 lx), it switches to CW circling [Fig. 1(c), (d)]. At intermediate intensities (~ 2,000 lx), cells actively switch handedness, transitioning from CW to CCW due to light adaptation (Fig. S3, Movie S4). Systematic quantification across different light intensities reveals that CCW trajectories dominate at 150 lx (~ 75%) and CW trajectories dominate above 10,000 lx (*>* 58%) [Fig. 1(e)]. Notably, RnT at 0–100 lx is characterized by stochastic switches between straight runs and abrupt turns, whereas RnT at 500–1,000 lx combines runs, turns, and both CCW and CW trajectories. By measuring *φ* over 90 seconds, we estimated circling periods of ~ 30 s for CCW and ~ 12 s for CW swimming (see Supplementary Section 2.3). These results indicate that light intensity not only determines trajectory handedness, but also influences the radius and timescale of the circling, pointing to a light-intensity dependent symmetry breaking mechanism in sensor-actuator responses.

### Sensor-actuator coupling

To understand the mechanism, we track the cell’s eyespot and flagella under different light conditions. Free-swimming cells were tracked at 1,000 fps, to resolve the complete flagellar waveform whenever the beat plane was perpendicular to the light direction during cell rolling (i.e., *ψ* = *π/* 4 or 5*π/* 4). We compared flagellar beats when the eyespot directly faced the light (light phase) versus when it was shaded (dark phase), quantifying beat patterns by tracking flagellar outlines and extracting elliptical orbits (*N* = 12 cells, ~ 20 frames per beat cycle) (see Supplemental Section 2.5).

During CCW swimming at low light intensities (~ 150 lx), the *cis-* flagellum exhibited a more extended waveform than the *trans-* flagellum (denoted by Δ*h >* 0), with a different amount of modulation in the light and dark phases (*P <* 0.01) [Fig. 2(a)]. This asymmetry in beat extension results in unequal turning during the light and dark phases and *cis-* flagellum dominance. This tunable flagellar dominance subsequently modulates cellular dynamics over longer timescales (~ 0.5 s) via the coupling between the eyespot and cell rolling. To quantify the relationship between eyespot position (*ψ*) and cell orientation (*φ*), we define sin(*δ*) to represent the phase of helical swimming, obtained by: *φ* = sin(*δ*)(*φ* _*max*_*− φ* _*min*_)*/* 2 + (*φ* _*max*_ + *φ* _*min*_)*/* 2. Here, *φ* _*max*_ and *φ* _*min*_ represent the maximum and minimum values of *φ* in each rolling cycle. When sin(*ψ*) increases from −1 to 1, the eyespot detects light effectively (light phase), while a decreasing sin(*ψ*) corresponds to the dark phase. During CCW swimming, sin(*δ*) and sin(*ψ*) were synchronized [Fig. 2(b)], indicating that the cells turned in the CCW direction (*φ* increases) during the light phase and in the CW direction (*φ* decreases) during the dark phase. We measured the average turning angle after one complete beat cycle Δ*φ* (see Supplemental Section 2.4), and obtained Δ*φ* = 3.96°± 0.91° (mean sem, used throughout unless otherwise stated) in the light phase and Δ*φ* = *−* 1.89°± 0.31° in the dark phase. The accumulative change of Δ*φ* over the entire rolling cycle ultimately generates a CCW circular trajectory with a relatively larger radius (~ 0.24 *±* 0.08 mm) [Fig. 2(c)].

**FIG. 2.**
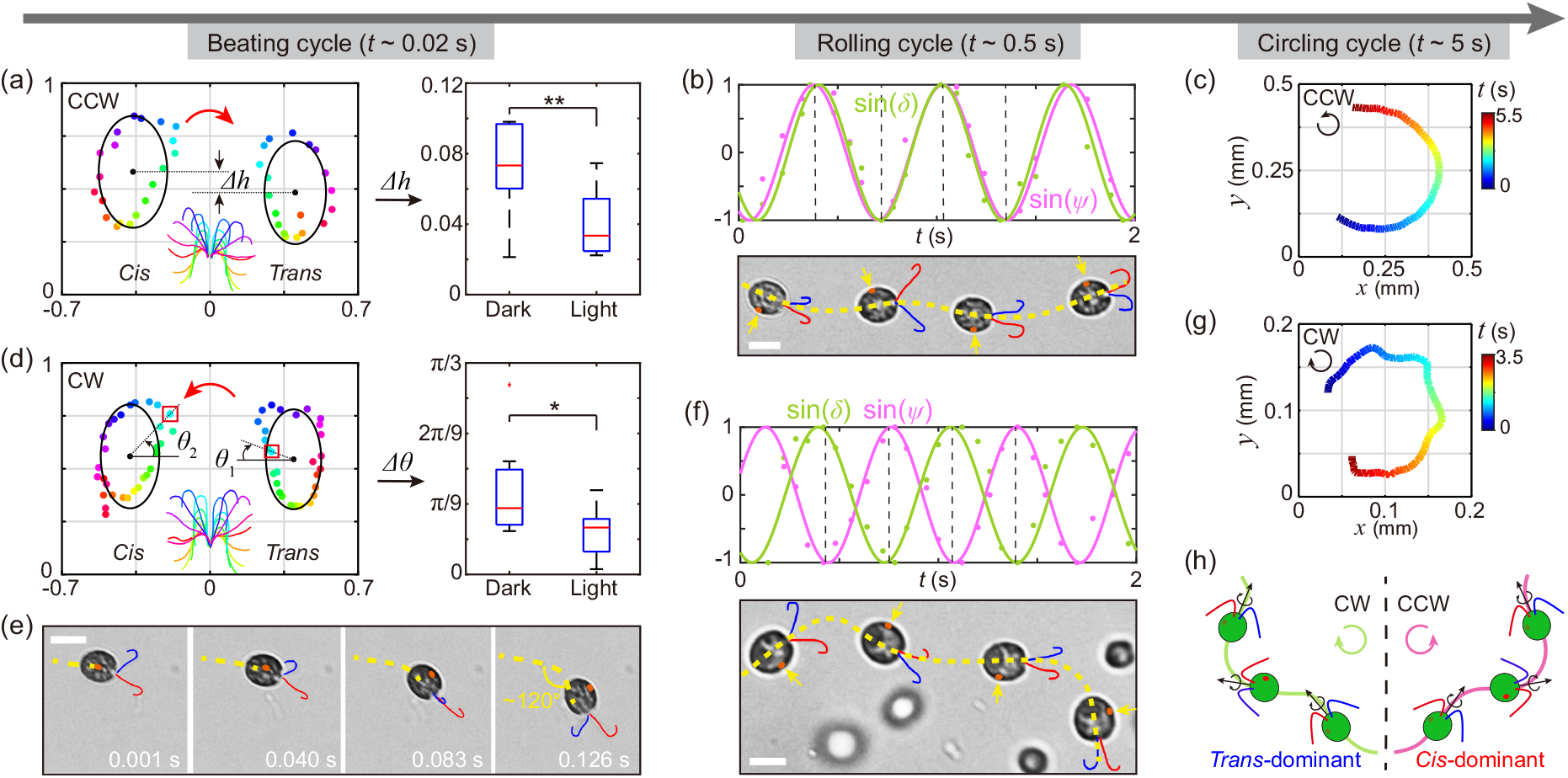
Live-cell experiments reveal the multiscale nature of sensor-actuator coupling and flagellar dominance. (a, d) Flagellar orbits (normalized by cell body length) and waveforms over a beat cycle for CCW (a) and CW (d) circling. Colored dots and waveforms indicate time-ordered frames (*N* =12 cells); red arrows denote the cell’s turning direction. (b, f) Comparison of sin(*ψ*) and sin(*δ*) phases, with overlaid cell images for CCW (b) and CW (f) circling. Dashed lines mark timepoints shown in cell images. (c, g) Example trajectories of CCW and CW circling (Movies S5 and S6). (e) Time-lapse images show the sharp turn during CW swimming (Movie S7). *Cis-* and *trans-* flagella are colored red and blue, respectively, eyespot highlighted in orange. (h) Schematic representation of how opposing flagellar dominance leads to CW or CCW circling. Scale bars: 10*µ* m.

Conversely, at high light intensity (~ 15,000 lx), no significant difference in beat extension between the flagella (Δ*h ~* 0) was observed for CW swimming (see Supplementary Section 2.5). Instead, a phase difference emerged between the two flagella (Δ*θ* = *θ* _2_ *−θ* _1_, *P <* 0.05) [Fig.2(d)], flipping flagellar dominance so that the *trans-* flagellum led CW turning. Notably, sharp turning events (up to 60° within 0.1 s) occurred when the eyespot directly faced the light (sin(*ψ*) *≈* 0) [Fig. 2(e)], where an abrupt reorientation of the flagellar beat plane was observed (Movie S5). This process induces a phase shift between the eyespot and helical swimming trajectory [Fig. 2(f)], so that during CW swimming, the eye-spot is located on the opposite side compared to CCW swimming, and the cell turns sharply in the light phase, generating smaller radii (~ 0.07*±* 0.02 mm) and flower-like trajectories [Fig. 2(g)].

Hence, light-induced subcellular asymmetries in beat extension, phase, and beat plane orientation regulate the coupling between eyespot position and flagellar dominance. While beat extension and phase can be measured by tracking flagella beats of free-swimming cells [Fig. 2(a), (d)], the change in beat plane orientation cannot be quantified easily due to cell rolling. In the following, we further elucidate how the beat plane orientation is connected to the orthogonal circular swimming by micropipette aspiration experiments and hydrodynamic modeling.

### Orthogonal swimming

Individual *Chlamydomonas* cells were gently aspirated and properly oriented to allow top-down high-speed imaging (3,000 fps) of both flagella [32]. Blue light (470 nm) pulses (2 Hz, close to the cell’s natural rolling frequency) were applied at different intensities using an optical fiber coupled to an LED (see more details in Supplementary Section 2.7). Under high-intensity light (~ 15,000 lx), both *cis-* and *trans-* flagella beat planes shifted markedly toward the eyespot, with a maximum reorientation of about 0.3 radians [Fig. 3(a), (b), Movie S8]. In contrast, at low intensity (~ 50 lx), both flagella directed away from the eyespot with a smaller shift of 0.1-0.2 radians [Fig. 3(c), (d), Movie S9]. After light removal, the beat planes returned to their original positions.

**FIG. 3.**
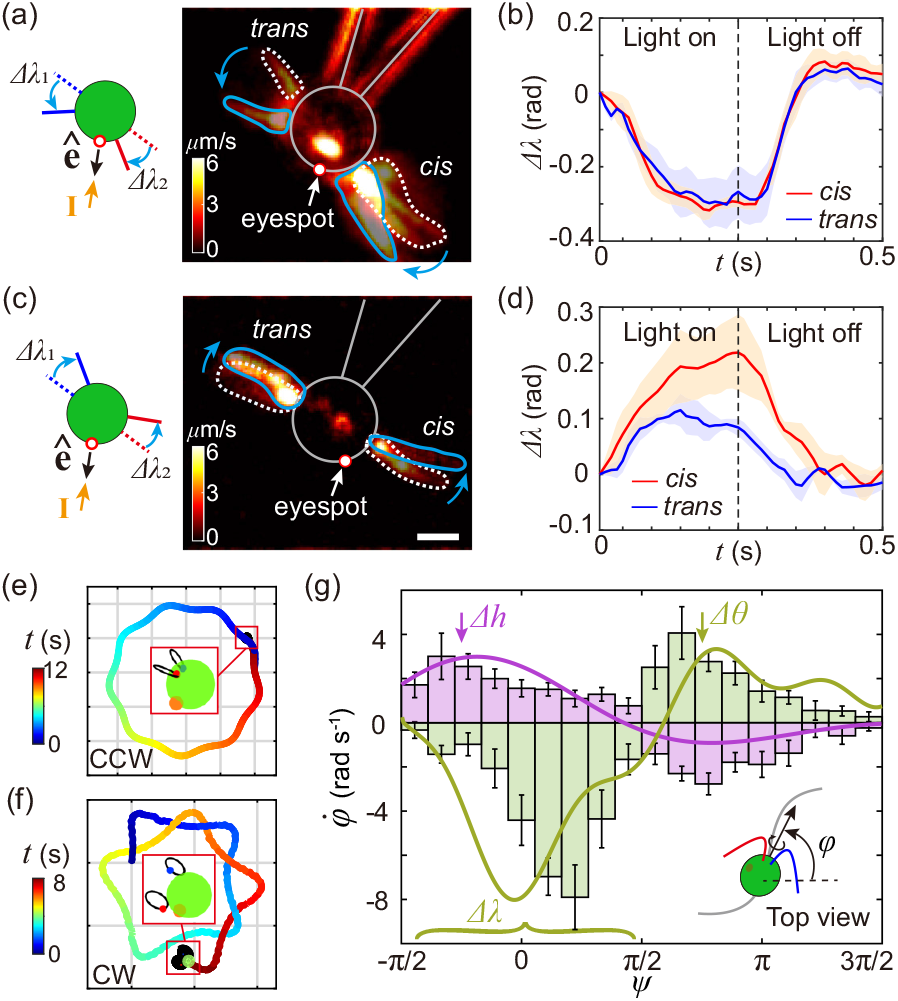
Beat plane reorientation enables circling behavior in *Chlamydomonas*. (a-d) Optical flow tracking shows beat plane reorientation toward the eyespot under high light (a, b) and away under low light (c, d). White dashed line: pre-illumination beat plane; solid blue line: post-illumination. Δ*λ* indicates the angular change of the beat plane before and after light stimulation. Shaded regions in (b,d) show the SEM (*N* = 5 measurements for b, *N* = 3 for d). (e, f) Simulated CCW (e) and CW (f) trajectories reconstructed from experimental data in (b, d) (Movies S10 and S11). Grid size in (e): 50 *µ* m, (f): 100 *µ* m. (g) Coupling between 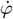 and *ψ*; green (CW) and purple (CCW) curves show simulation results, and bars show experiments (*N* = 8 per case). Scale bar: 5*µ* m

The mechanism of light-induced orthogonal circular swimming is distinct from normal phototaxis where cells swim parallel to the incident light. The two flagella of *Chlamydomonas* are asymmetrically positioned relative to the eyespot (offset by *π/* 4) [24]. When the flagella exert unbalanced forces due to photoresponses, this inherent structural asymmetry helps generate unbalanced torques perpendicular to the light direction, resulting in phototactic steering of the cell parallel to the light stimulus. Yet, orthogonal circular swimming is non-trivial as the cell has to achieve zero torques along all other directions except the light direction normal to the swimming plane. Next we use hydrodynamic modeling to demonstrate that light-dependent beat plane reorientation (as observed in our experiments) can indeed compensate for this asymmetry by redirecting the net force and torque along the light direction, enabling stable orthogonal circular swimming.

We revisit a three-sphere hydrodynamic model of *Chlamydomonas* [25, 32, 33]. This model can readily capture phototaxis where cells swim parallel to light. Here we extend this model to incorporate light-induced orthogonal swimming. As evidenced by the experiments, the cells can regulate their flagellar beat waveforms, force, and beat plane orientation in response to the light signal *S* (*t*), where *S* (*t*) is given by:

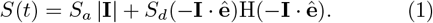

Here **I** and **ê** are the light vector and the eyespot vector, respectively, while *S* _*a*_ and *S*_*d*_ are the coupling constants for ambient and directional light. The Heaviside function (H) accounts for light shading. The flagella beat extensions (Δ*h*) and beat phases (Δ*θ*) are modulated by: *h*_*i*_ = *h*_0_ + *K*_*h,i*_ log(*S* (*t*)) and *c*_*i*_ = *c*_0_ + *K*_*c,i*_ log(*S* (*t*)) in CCW and CW modes respectively (*i* = 1, 2 for *trans* - and *cis*-flagellum). *K*_*h,i*_ and *K*_*c,i*_ are light-dependent coupling constants (see Supplementary Section 3.1-3.3 for model details).

Model simulations reproduced the salient features of orthogonal CCW and CW trajectories, with negligible displacement along the light direction [Fig. 3(e), (f)] (Supplementary Section 3.4-3.8). Trajectory handedness is determined by the sign of the net torque generated by the two flagella, which switches when the beat planes reorient. The curvature of the circular trajectory is proportional to the magnitude of the beat plane reorientation. We compared simulated reorientation rates in the orthogonal plane 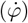 with experimental measurements in free-swimming cells [Fig. 3(g)]. Three distinct modes of curvature modulation were evident: (1) When *ψ* = *−π/* 4 (light phase), spatial asymmetries in beat extension (Δ*h*) between flagella drive CCW turning; (2) when *ψ* = 3*π/* 4 (dark phase), temporal phase asymmetries (Δ*θ*) drive CW turning; (3) at *ψ* = 0, when the eyespot directly faces light, beat plane reorientation reaches its maximum. Under intense light (large Δ*λ*), this triggers sharp turns (corresponding to large 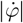) and flower-like curvature in CW trajectories, whereas under low light, smaller Δ*λ* leads to smoother trajectories. Thus, the model quantitatively captures how beat plane reorientation finely controls trajectory curvature and handedness, in agreement with experimental data in both free-swimming and pipette-held cells.

### Navigation strategies

Finally, we explored the ecological relevance of light-induced circular swimming by examining how *Chlamydomonas* utilizes curvature modulation for navigation in spatially structured light environments. Using a digital micromirror device (DMD), we generated two types of spatial light fields and tracked single-cell trajectories over five minutes [Fig. 4(a), (b)].

**FIG. 4.**
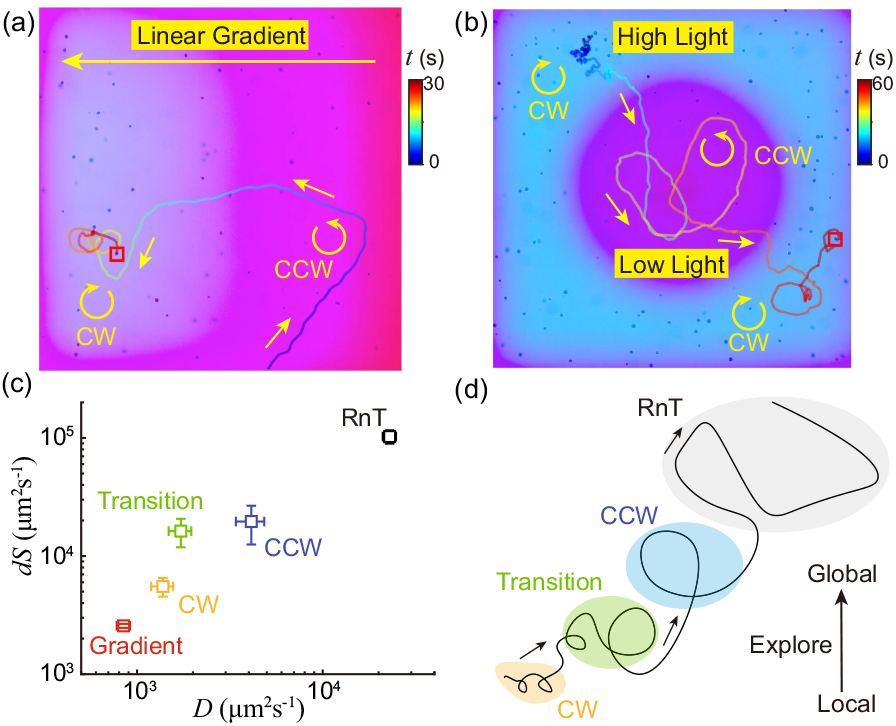
Adaptive navigation in structured light fields. (a) In a spatial light gradient, cells alternate between CW and CCW circling, navigating toward brighter regions (Movie S12). (b) In a stepped light field, cells initially swim CW, then switch to larger-radius CCW circling before re-entering high-light areas (Movie S13). (c) Comparison of diffusion constant (*D*) and exploration rate (*dS*) across different navigation strategies (*N* = 6 per case) (“Gradient” refers to trajectories in the light gradient where different modes are mixed). (d) Schematic of the robust navigation strategy exhibited by *Chlamydomonas* cells as they switch between CW, transition, CCW, and RnT behaviors, supporting both local and global exploration.

In a linear light gradient [Fig. 4(a)], cells entering low-light regions adopted a CCW circling “search mode” with a relatively large radius, effectively performing gradient ascent. Upon reaching high-light regions, cells switched to CW circling with a smaller radius, remaining localized in these areas for extended periods (*>* 60 seconds). In a step-gradient environment with a low intensity (~ 150 lx) in the center and a high intensity (~ 15,000 lx) in the periphery [Fig. 4(b)], cells in high-light zones exhibited noisy, CW circling behavior. When cells exited high-light regions, they immediately switched to the CCW search mode until re-entering high-light regions within a few cycles. These behaviors demonstrate that *Chlamydomonas* dynamically modulates trajectory curvature and handedness to flexibly explore spatially varying light fields, enabling strategies such as gradient ascent and re-entry into high-light regions.

To further quantify these navigation strategies, we calculated the long-time mean squared displacement (MSD) and area covered per unit time (*dS*) under uniform light field at different intensities [34] (see Supplementary Section 2.8 for definitions) [Fig.4(c)]. In darkness (~ 0 lx), cells exhibited RnT with the largest generalized diffusion coefficients (*D* = 22, 950 *±* 1, 882 *µm* ^2^*/s*), exploring wide areas (*dS* = 102, 667*±* 13, 167 *µm* ^2^*/s*). In low light (~ 150 lx), CCW swimming emerges with decreased diffusion (*D* = 4, 125 *±* 719 *µm* ^2^*/s*; *dS* = 19, 600 *±* 7, 067 *µm* ^2^*/s*). In high light (~ 15,000 lx), cells switched to CW swimming with further reduced diffusion (*D* = 1, 378 *±* 187 *µm* ^2^*/s*; *dS* = 5, 533 *±* 1, 000 *µm* ^2^*/s*).

In intermediate light intensities (~ 2,000 lx), where cells transition between CW and CCW modes, the diffusion is intermediate (*D* = 1, 720 *±* 240 *µm* ^2^*/s, dS* = 16, 267 *±* 4, 400 *µm* ^2^*/s*). In spatial gradients, with frequent mode switching and localized circling, diffusion was the lowest (*D* = 846 *±* 67 *µm* ^2^*/s*; *dS* = 2, 567 *±* 100 *µm* ^2^*/s*). Thus, by tuning trajectory curvature in response to the local light field, *Chlamydomonas* can select between localized search (small circles, low diffusion), global exploration (large circles, high diffusion) and more diffusive RnT, allowing efficient navigation in heterogeneous environments [Fig. 4(d)].

## Conclusions

In summary, we identify a novel subcellular symmetry-breaking mechanism of sensor-actuator response to light in *Chlamydomonas*. In particular, cells exhibit circular swimming behaviors with an intensity-dependent handedness in response to orthogonal illumination. Using a combination of live cell high-speed tracking, micropipette aspiration assays, and hydrodynamic modeling, we show that circular swimming is triggered by the modulation of flagellar dominance, through specific changes in waveform, phase and beat plane orientation. Previous studies have demonstrated the necessity of non-planar flagellar beating for helical swimming [32], here we further show that the reorientation of flagellar beat plane is crucial for the transition from parallel phototaxis to orthogonal photoresponses. This additional degree of freedom offers the cell with a high degree of flexibility to robustly navigate in three dimensions via helical kilotaxis. Transitions from helical swimming to circular swimming in response to environmental cues have been observed in many eukaryotic species such as spermatozoa, diatoms, and euglenids [12, 35–38], despite having different motility actuation mechanisms, underscoring trajectory curvature modulation as a fundamental navigation strategy in microswimmers.

Finally, we remark that while helical klinotaxis is universal for biological microswimmers, the implementation differs across species. For example, sperm cells exhibit chemotaxis by coupling gradual trajectory drifts with sharp turns, causing their circling centers to shift over times and approach chemical sources [36]. *Euglena gracilis* transition between swimming modes with different diffusion constants to perform light gradient ascent or descent [38, 39]. Here, we present another navigation strategy found in *Chlamydonomas reinhardtii*, where cells perform light gradient ascent by transitioning between large-radius circles in low-light regions and small-radius circles in high-light regions, effectively tuning the diffusion constants and exploration area. Identifying deeper connections between these different mechanisms may uncover general biophysical laws for microswimmer navigation. Taken together, our findings identify the role of active flagella beat plane reorientation in cell motility and mechanical torque balance for complex stimulus-dependent maneuvers, as a general mechanism for curvature control in microswimmer navigation.

## ACKNOWLEDGMENTS

This work was supported by the Research Grants Council of Hong Kong through the General Research Fund (No. 27208421 and No. 17303423) and the Croucher Foundation (A.C.H.T.). This work was also funded by the European Research Council (ERC) under the European Union’s Horizon 2020 research and innovation programme grant No. 853560 EvoMotion, and a Springboard Award from the Academy of Medical Sciences and Global Challenges Research Fund (SBF003\1160) (K.Y.W.).

## Notes

### Competing Interest Statement

The authors have declared no competing interest.

