## Supplementary Information for "Light-dependent switching of circling handedness in microswimmer navigation"

#### 1 Supplementary movies

**Movie S1**: Run-and-tumble trajectory of *Chlamydomonas* in darkness (∼0 lx). The video was recorded at 10 fps and played back at 3× speed.

**Movie S2**: Counterclockwise (CCW) swimming trajectory of *Chlamydomonas* under low light intensity (∼150 lx). The video was recorded at 10 fps and played back at 3× speed.

**Movie S3**: CW swimmng trajectory of *Chlamydomonas* under high light intensity (∼15000 lx). The video was recorded at 10 fps and played back at 3× speed.

**Movie S4**: Transition from CCW to clockwise (CW) trajectory of *Chlamydomonas* at intermediate light intensity (∼2000 lx). The video was recorded at 10 fps and played back at 3× speed.

**Movie S5**: High-resolution recording of a detailed CCW swimming trajectory of *Chlamydomonas*. The video was captured at 1000 fps and replayed at 0.2× the original speed.

**Movie S6**: High-resolution recording of a detailed CW swimming trajectory of *Chlamydomonas*. The video was recorded at 1000 fps and played back at 0.2× the original speed.

**Movie S7**: Sudden turning of *Chlamydomonas* under high light intensity when the eyespot directly faces the light. The video was recorded at 1000 fps and played back at 0.02× the original speed.

**Movie S8**: Reorientation of the beat plane towards the eyespot in *Chlamydomonas* under high light stimulation. The video was recorded at 3000 fps and played back at 1/60× the original speed.

**Movie S9**: Reorientation of the beat plane away from the eyespot in *Chlamydomonas* under low light stimulation. The video was recorded at 3000 fps and played back at 1/60× the original speed.

**Movie S10**: Simulation demonstrating the smooth CCW trajectory of *Chlamydomonas*.

**Movie S11**: Simulation demonstrating the flower-like CW trajectory of *Chlamydomonas*.

**Movie S12**: *Chlamydomonas* performing gradient ascent in a linear gradient light field. The video was recorded at 10 fps and played back at 2× speed.

**Movie S13**: *Chlamydomonas* switching between CCW and CW modes to re-enter a bright region in a stepped light field. The video was recorded at 10 fps and played back at 2× speed.

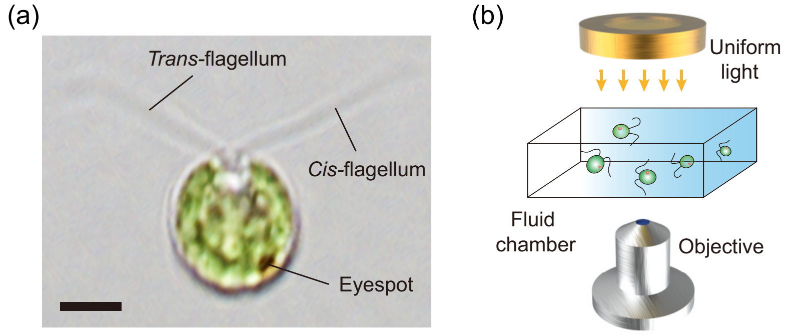


Figure S1: (a) Image of *Chlamydomonas reinhardtii*. Scale bars, 5 $\mu$m. (b) Schematic of the experimental set-up.

### 2 Experimental analysis

#### 2.1 Cell culture and experiment set-up

The wild-type *Chlamydomonas reinhardtii* (NIES-2238, 137C $mt^+$) was cultured in TAP medium at ∼24 °C under a 12:12 light/dark cycle (3000 lx). Cells in exponential phase (2–3 days) were used for experiments.

Free-swimming *Chlamydomonas* cells were observed in custom fluid chambers [2 cm × 2 cm × 100 $\mu$m; (Fig. S1)] assembled with tapes between a glass slide and a cover slip. The cells were imaged using an inverted microscope (Nikon Eclipse Ti2). Chamber depth was measured by caliper. We also performed control experiments in 300 $\mu$m-deep chambers and obtained similar results, indicating boundary effects are negligible.

Uniform illumination was provided by a microscope lamp. For dark conditions, a long-pass filter (Thorlabs FGL630M) blocked stimulating light, allowing only red background illumination which cells do not respond. Cell motion was recorded at 10 fps (Basler acA2040-90uc); flagellar beats were imaged at ≥1000 fps (Phantom V2012).

Structured light fields [e.g., gradients in main text Fig. 4(a)] were generated using a DMD (Mightex Polygon 1000) by dividing the field of view into stripes of varying brightness ($n > 10$) via rapid temporal modulation (1000 Hz). Blue light (∼ 470 nm) from the DMD stimulated *Chlamydomonas*; a red filter provided red background light.

#### 2.2 Trajectory reconstruction

We used both automatic and manual tracking for *Chlamydomonas* trajectories. For high-throughput analysis [e.g., trajectory handedness in main text Fig. 1(e)], automatic tracking was performed with TrackMate (ImageJ), with further analysis via custom scripts.

For long-time tracking experiments of individual cell's handedness and changes in curvature due to adaptation [e.g., main text Fig. 1(b)–(d)], manual tracking was used. Cells were followed for up to 5 min by adjusting the microscope stage, with relative coordinates corrected using immobile references to reconstruct absolute trajectories (Fig. S2). This allowed for continuous tracking beyond the FOV limits. Combining both methods enabled efficient statistical analysis and detailed single-cell dynamics.

To determine the fractions of behavioral states across light intensities [e.g., main text Fig. 1(e)], we analyzed trajectories of ∼ 130 cells per light intensity (15 intensities, >2,000 cells in total) using TrackMate in ImageJ. Behavioral states (run-and-tumble, CCW, CW) were classified with custom scripts: short or immobile tracks were excluded, remaining trajectories were categorized based on curvature, and all classifications were manually verified for accuracy.

#### 2.3 Orientation comparison

In addition to the three major trajectory types shown in main text of Fig. 1, at intermediate light (∼ 2000 lx), *Chlamydomonas* displays a transition behavior and switches from CW to CCW over times [Fig. S3(a)-(d)]. We quantified trajectory handedness and curvature by calculating the orientation angle $\varphi$ (relative to the lab frame), which increases for CCW and decreases for CW circling [Fig. S3(e)-(f)].

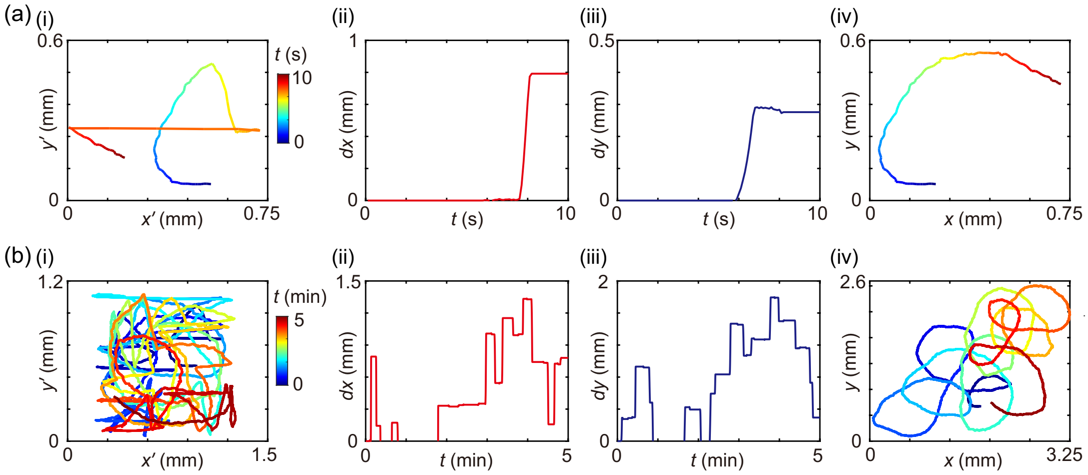

Figure S2: Two-step transformation from relative to absolute coordinates for short (a) and long (b) trajectories. (i) cell trajectory in moving FOV; (ii–iii) stage displacement in $x$ and $y$; (iv) reconstructed lab-frame trajectory.

Further analysis of the orientation rate ($\dot{\varphi}$) revealed that CW trajectories have broader, time-varying distributions, while CCW are more constant [Fig. S3(g)-(h)]. This time-varied change in CW indicated increasing trajectory radius, likely caused by light adaptation [1].

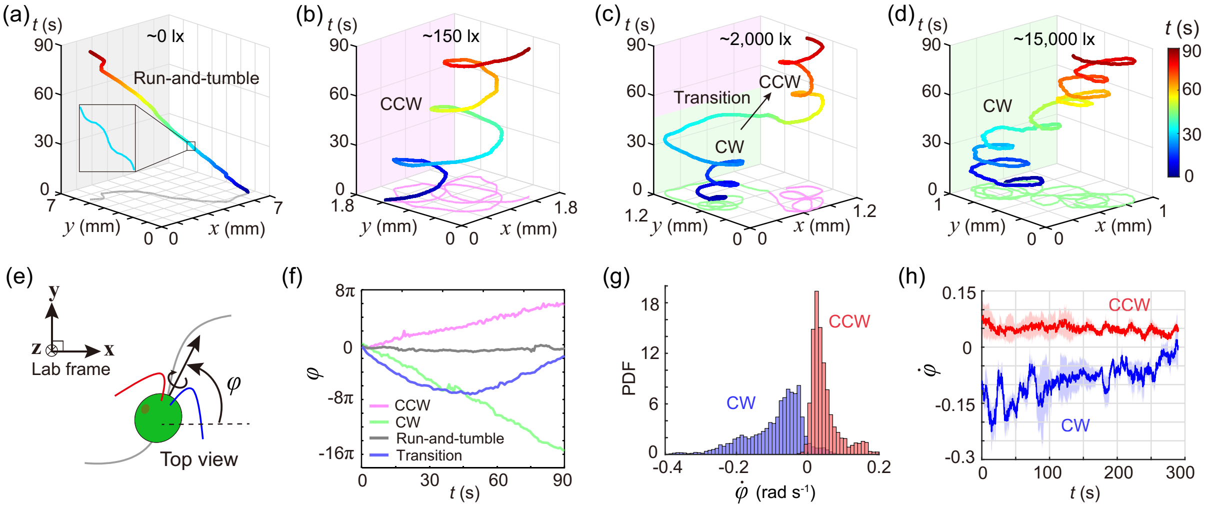

Figure S3: *Chlamydomonas* swimming modes: (a) run-and-tumble, (b) CCW circle, (c) CW-to-CCW transition, (d) CW circle. (e) Definition of orientation angle $\varphi$. (f) Accumulated $\varphi$ change over 90 s for each mode. (g) PDF of orientation rate $\dot{\varphi}$. (h) Temporal distribution of $\dot{\varphi}$ in CCW and CW motion ($N = 4$ cells each).

#### 2.4 Flagella dominance

Here we briefly discussed the transition of flagella dominance as shown in Fig. 2 of the main text. Flagellar dominance is determined by analyzing the turning direction of *Chlamydomonas* after each flagellar beat during light/dark phases (beating plane perpendicular to light). For CCW swimming, the *cis*-flagellum beats more strongly, generating positive turning angles ($\Delta\varphi$) in light phases and negative angles in dark phases [Fig. S4(a) and (c)], resulting in sustained CCW circling (mean $\Delta\varphi_{\mathrm{light}} = 3.96 \pm 0.91°$, $\Delta\varphi_{\mathrm{dark}} = -1.89 \pm 0.31°$, $n = 10$; [Fig. S4(d)].

For CW swimming, the *trans*-flagellum dominates, with negative $\Delta\varphi$ in light phases and positive $\Delta\varphi$ in dark phases [$\Delta\varphi_{\mathrm{light}} = -0.94 \pm 0.29°$, $\Delta\varphi_{\mathrm{dark}} = 2.53 \pm 0.37°$, $n = 9$; Fig. S4(b), (e) and (f)]. While the average $\Delta\varphi$ remains positive, additional beat modes (e.g., beating plane reorientation) explain the observed CW circling. Thus, flagellar dominance correlates with turning direction, modulated by flagellar beat extension and beat phase.

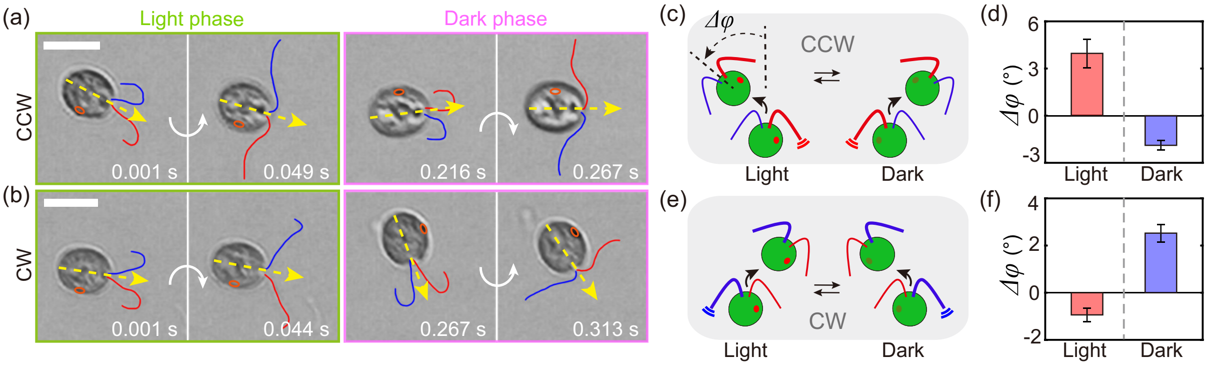


Figure S4: *Chlamydomonas* exhibits switchable flagella dominance. (a,b) Time-lapse sequences for CCW and CW circling; white arrows show the cell's turning direction. (c,e) Schematics of flagella dominance and (d,f) average orientation change per beat ($\Delta\varphi$) for CCW ($N = 10$) and CW ($N = 9$) cells, respectively. *cis-* and *trans*-flagella are denoted in red and blue; eyespot is denoted in orange. Scale bars: 10 $\mu$m.

#### 2.5 Flagella tracking

To analyze flagellar beat patterns, we manually tracked $n = 6$ *Chlamydomonas* cells swimming in both CCW and CW modes, focusing on periods when the beat plane was parallel to the imaging plane when the eyespot faced toward or was shaded from the light. Flagellar outlines were digitized at high temporal resolution (~500 points per frame, 20 frames per beat cycle at 1000 fps), and the mean coordinates of the *cis-* and *trans*-flagella were used to fit ellipsoidal orbits [Fig. S5a]. We extracted key geometric parameters ($a_i$, $b_i$, $h_i$, $l_i$, $\theta_i$) to quantify the size and position of the flagellar beat orbits, enabling direct comparison of beat patterns in different swimming modes.

During CCW swimming, the *cis*-flagellum exhibited significantly greater beat extension ($\Delta h > 0$), especially under the dark phase ($P < 0.01$), while the phase difference between the flagella ($\Delta\theta$) showed no significant change [Fig. S5(b)]. In contrast, CW swimming was characterized by light-induced asymmetry in the phase difference ($P < 0.05$), rather than differences in beat extension [Fig. S5(c)]. These results indicate that CCW and CW swimming rely on distinct mechanisms for modulating flagella dominance: beat extension for CCW and phase shift for CW.

In addition to beat extension and phase difference, we identified a previously unreported beating mode during CW swimming: when the eyespot directly faces the light, the cell makes rapid, large turns (up to 60° in 0.1 s) with both flagella bending to the same side [0.083 s in Fig. S6(a)]. This cannot be explained by in-plane beat changes alone. Comparison of experimental images and geometric reconstructions suggests these abrupt turns result from light-induced reorientation of the flagellar beat planes [Fig. S6(b), (c)], which is further supported by the micropipette experiments shown in the main text Fig. 3(a).

#### 2.6 Phase coupling relationships

To describe the phases of cell orientation and eyespot position, we used $\sin(\delta)$ and $\sin(\psi)$ to reveal distinct phase coupling for CCW and CW swimming. The cell orientation ($\varphi$) was directly tracked [Fig. S7(a)], and $\sin(\delta)$ was calculated as: $\varphi = \sin(\delta)(\varphi_{max} - \varphi_{min})/2 + (\varphi_{max} + \varphi_{min})/2$, where $\varphi_{max}$ and $\varphi_{min}$ are the maximum and minimum orientations in each rolling cycle.

The eyespot position during cell rolling ($\psi$) could not be measured directly. Instead, we estimated $\sin(\psi)$ by $\sin(\psi) = l_e/l_b$ [Fig. S7(b)], where $l_b$ is the cell radius and $l_e$ is the distance from the eyespot to the cell's long axis. As the cell rotates, $l_e$ oscillates between $-l_b$ and $+l_b$. By measuring the length of $l_e$ over time [Fig. S7(c)], we obtained the value of $\sin(\psi)$.

#### 2.7 Micropipette experiment

In micropipette experiments, individual cells were gently immobilized at the tip of finely polished micropipettes, and their orientation was precisely adjusted with a micromanipulator to allow top-down observation of the flagella [Fig. S8(a) and (e)]. High-speed videos were recorded at 3000 fps to capture rapid flagellar dynamics.

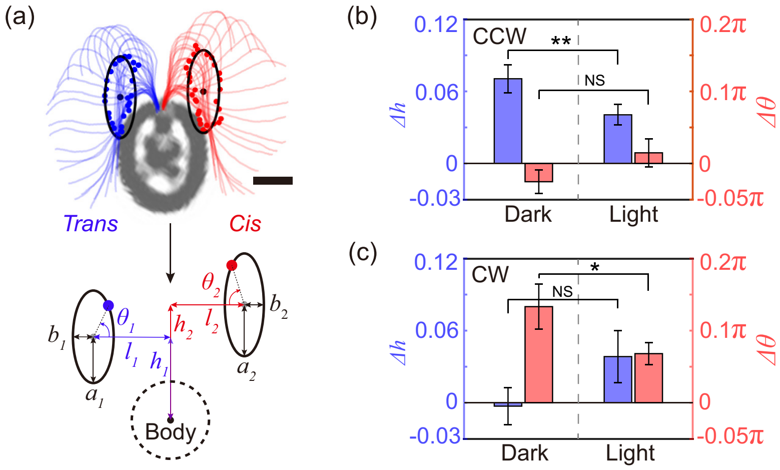


Figure S5: (a) Flagellar waveforms were tracked frame by frame and averaged coordinates were fitted as a pair of elliptical orbits. Orbit size is described by $a_i$ and $b_i$, position by $h_i$ and $l_i$, and beat phase in a cycle by $\theta_i$ ($i = 1, 2$ for *trans*- and *cis*-flagellum). Scale bars: 5 $\mu$m. (b,c) Differences in flagella extension ($\Delta h$) and phase ($\Delta\theta$) during CCW (b) and CW (c) swimming.

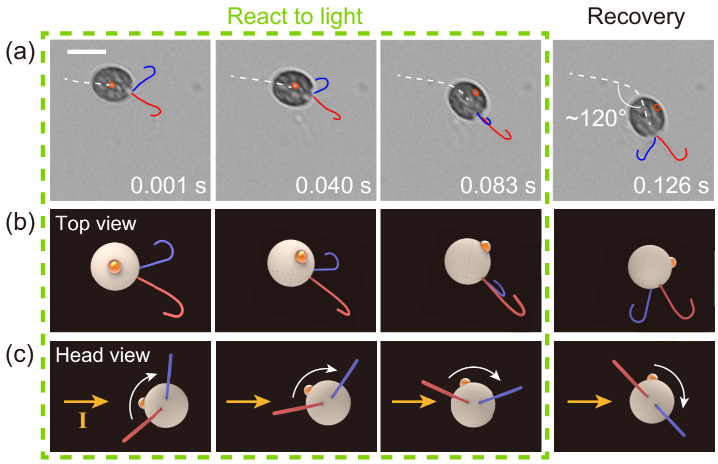


Figure S6: *Chlamydomonas* exhibits sharp turning during CW swimming when its eyespot detects light directly. (a) Time-lapse sequence showing sharp turning; *Cis*- and *trans*-flagella are marked in red and blue, with the eyespot highlighted in orange. (b) Top view and (c) head view comparing the flagellar beat plane geometry during swimming. Scale bars, 10 $\mu$m.

We analyzed the videos using custom Matlab scripts based on the Horn–Schunck optical flow algorithm [2–4]. Videos were first converted to grayscale and partitioned into cycles of 50 frames, corresponding to a full beat cycle. Background images were generated by averaging all frames in a cycle and then subtracted from each frame. Optical flow was then calculated to estimate the spatial domain of flagellar beating. The mean optical flow per cycle was visualized as a heatmap, from which we estimated the beat plane's orientation and angle relative to the eyespot.

Cells were exposed to 2 Hz square-wave blue light at high ( 15,000 lx) or low ( 50 lx) intensities, mimicking natural periodic signals experienced by a rolling cell. Before illumination, the angle between flagellar beat planes was about $\pi$ [Fig. S8(b) and (f)]. Light stimulation caused the planes to reorient [Fig. S8(c) and (g)]. Overlayed optical flow maps showed that high-intensity light reoriented beat planes toward the eyespot, while low-intensity light reoriented them away [Fig. S8(d) and (h)]. After the stimulus, planes gradually returned to their original orientations.

To quantitatively characterize these dynamics, we tracked changes in beat plane for each flagellum over 10 cycles at both light intensities. Example time courses of the light pulse and the angles between each flagellar beat plane and the eyespot ($\lambda_1$ and $\lambda_2$) are shown in Fig. S9(a) (high light) and Fig. S10(a) (low light). Data from each cycle were averaged to yield the mean temporal evolution for *cis*- and *trans*-flagella [Fig. S9(b) and (c) and Fig. S10(b) and (c)]. Multiple measurements ($N = 5$ for high light,

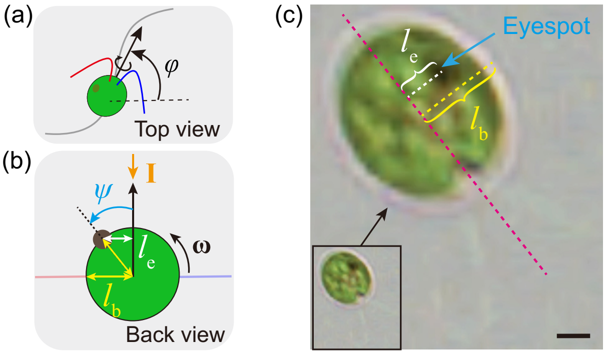


Figure S7: (a) Definition of cell orientation $\varphi$. (b) Definition of eyespot position $\psi$. $\sin(\psi) = l_e/l_b$. (c) Direct measurement of $l_e$ and $l_b$ from a real *Chlamydomonas* image. Scale bars, 10 $\mu$m.

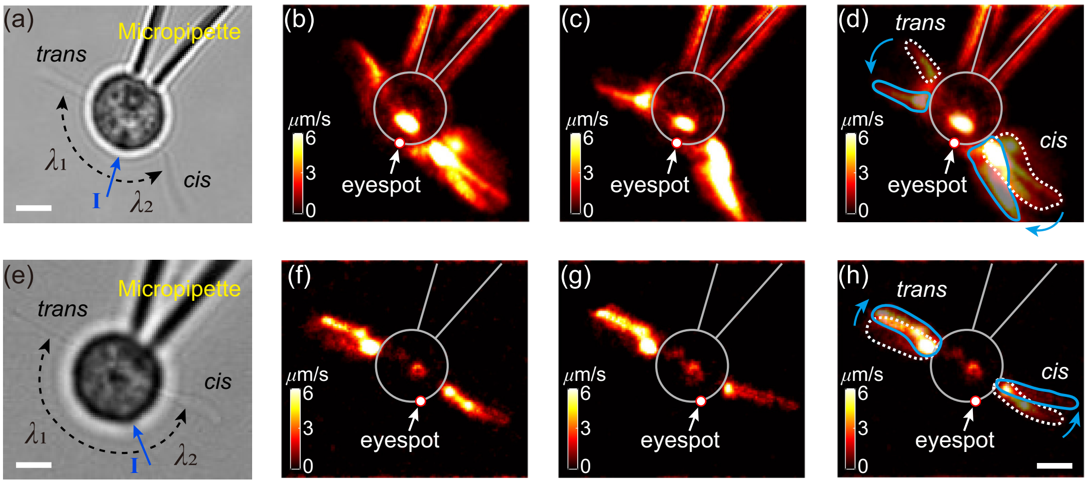


Figure S8: (a–d) Micropipette experiments under high-intensity light. (e–h) Micropipette experiments under low-intensity light. (a, e) Raw images of the experiments; (b, f) Optical flow maps before illumination; (c, g) Optical flow maps after illumination; (d, h) Overlays of the optical flow maps. Scale bars, 5 $\mu$m.

$N = 3$ for low light) showed consistent results (Fig. S11). Fitting these data revealed that the beat plane angles followed a half-cycle sinusoidal response to light, resulting in four fitted functions for the *cis*- and *trans*-flagella at each intensity.

In our simulations, the beat plane of both flagella were modulated by experimentally fitted sine functions whenever the eyespot detected light, ensuring the simulated dynamics matched experimental observations. This temporal profile was used in all subsequent simulations.

#### 2.8 MSD and explore rate ($dS$)

In the main text, we evaluated the navigation of *Chlamydomonas* using both the generalized diffusion coefficient and the convex hull area. The diffusion coefficient reflects how quickly the organism spreads spatially, while the convex hull area quantifies the total region explored.

Unlike simple Brownian motion, navigation in complex environments often displays anomalous diffusion [5], such as in cases of circular swimming (CW or CCW), or responses to gradients. In these situations, the mean squared displacement (MSD) deviates from the linear time dependence expected for normal diffusion, instead following a power-law relationship:

$$\mathrm{MSD}(t) = \left\langle (\mathbf{r}(t+\Delta t) - \mathbf{r}(t))^2 \right\rangle \propto (\Delta t)^{\alpha}, \tag{S1}$$

where $\mathbf{r}(t) = (x(t), y(t))$ is the position at time $t$, $\Delta t$ is the time lag, and $\langle \cdot \rangle$ denotes averaging over all time points. The exponent $\alpha$ characterizes the diffusion type: $\alpha = 1$ (normal diffusion), $\alpha < 1$ (subdiffusion), and $\alpha > 1$ (superdiffusion).

To make fair comparisons across different navigation strategies, we extract the generalized diffusion coefficient ($D_\alpha$) by fitting the MSD curve with [6, 7]:

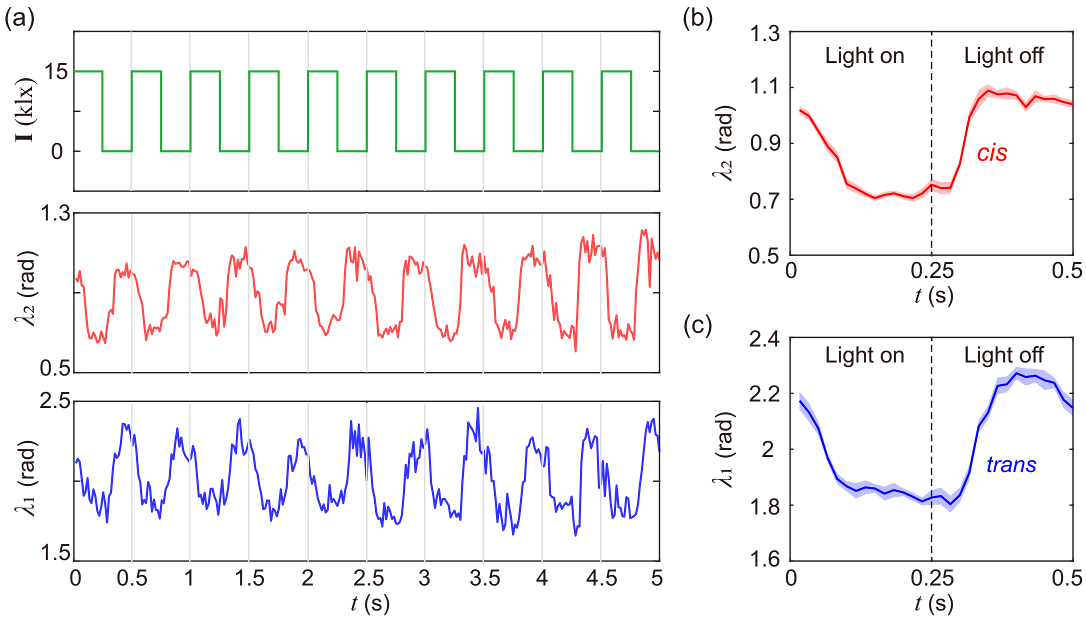


Figure S9: An example measurement of the micropipette experiment under high light intensity ($\sim 15,000$ lx). (a) Time courses of light intensity (I), the angle between the *cis*-flagellum beat plane and the eyespot ($\lambda_2$), and the angle between the *trans*-flagellum beat plane and the eyespot ($\lambda_1$) over 10 cycles of light stimulation. (b) Mean value of $\lambda_2$ over the 10 cycles. (c) Mean value of $\lambda_1$ over the 10 cycles. Shaded areas represent the standard error of the mean (SEM).

$$\mathrm{MSD}(t) = 4D_\alpha(\Delta t)^\alpha, \tag{S2}$$

where $D_\alpha$ and $\alpha$ were obtained by nonlinear regression on the log-log MSD curve at intermediate time scales to avoid artifacts from short-time ballistic motion and long-time confinement or boundary effects. This enabled quantitative comparison between normal and anomalous diffusion across different behavioral modes.

We also quantified exploration by calculating the convex hull area ($S$) of each trajectory and defining the exploration rate as $dS = S/t$, which captures how rapidly the cell explores new territory. Comparing trajectories of each mode revealed significant differences in the areas explored, reflecting a transition from global exploration to localized exploration (Fig. S12).

### 3 Hydrodynamic model

This section provides the details of the methods and simulation results of the hydrodynamic model. We first describe the model dynamics and numerical methods (Sections 3.1–3.3). We then systematically explore how varying the handedness and radius of orthogonal circular trajectories can be simulated and controlled (Sections 3.4–3.6). Finally, we compare simulation results to experimental data to investigate curvature modulation mechanisms in *Chlamydomonas* and summarize the main findings (Sections 3.7–3.8).

#### 3.1 Model dynamics

Our model is developed based on a previously established coarse-grained three-sphere representation [1, 4, 8]. In this model, the cell body is modeled as a large sphere ($\mathbf{r}_0$), while the two flagella are represented by smaller spheres ($\mathbf{r}_i$, $i = 1, 2$, corresponding to the *trans*- and *cis*-flagellum, respectively) [Fig. S13(a)]. The smaller spheres follow predefined elliptical orbits in planes $\pi_i$, simulating the effects of flagellar beats.

The motion of each flagellar sphere is composed of three parts: translation of the cell body ($\dot{\mathbf{r}}_0$), rotational motion of the cell body ($\Omega \times (\mathbf{r}_i - \mathbf{r}_0)$), and the movement of the sphere along its orbit ($\dot{\mathbf{r}}_i'$). Thus, the velocity of the flagellar spheres can be expressed as:

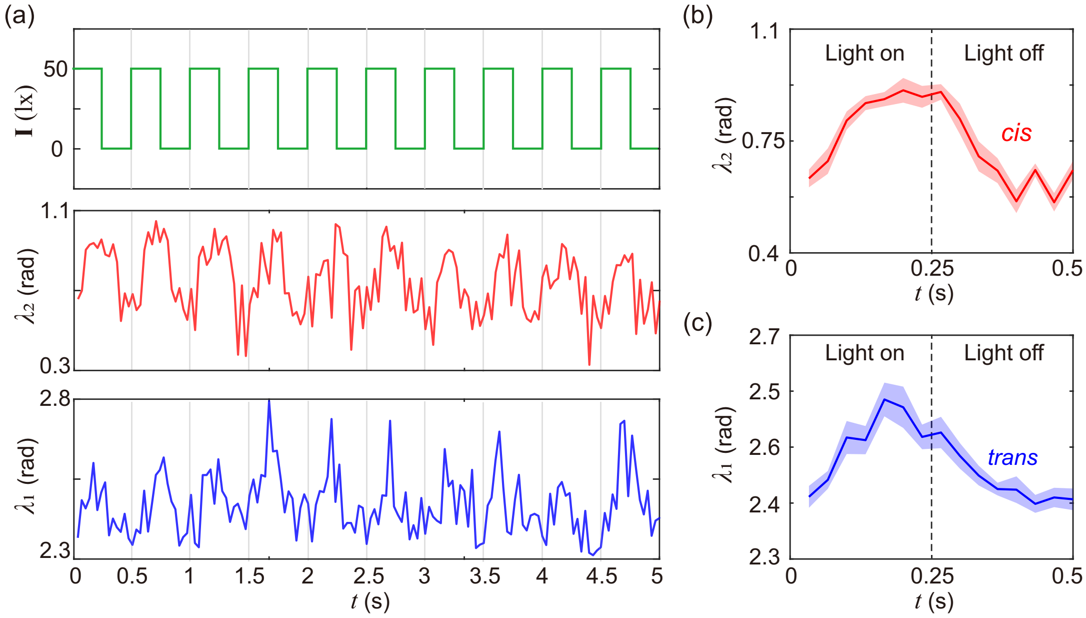


Figure S10: An example measurement of the micropipette experiment under low light intensity ($\sim 50lx$) (a) Time courses of light intensity (I), the angle between the *cis*-flagellum beat plane and the eyespot ($\lambda_2$), and the angle between the *trans*-flagellum beat plane and the eyespot ($\lambda_1$) over 10 cycles of light stimulation. (b) Mean value of $\lambda_2$ over the 10 cycles. (c) Mean value of $\lambda_1$ over the 10 cycles. Shaded areas represent the standard error of the mean (SEM).

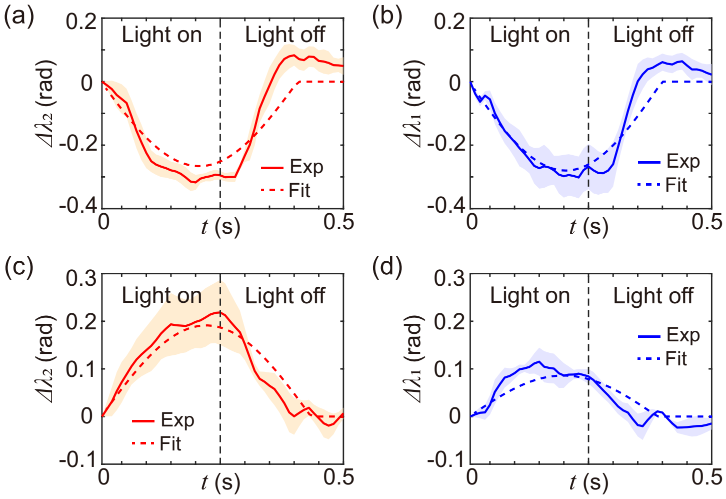


Figure S11: Fitted dynamics of beat plane angles under different light intensities. (a) *Cis*-flagellum ($\lambda_2$) under high light (~15,000 lx). (b) *Trans*-flagellum ($\lambda_1$) under high light. (c) *Cis*-flagellum under low light (~50 lx). (d) *Trans*-flagellum under low light.

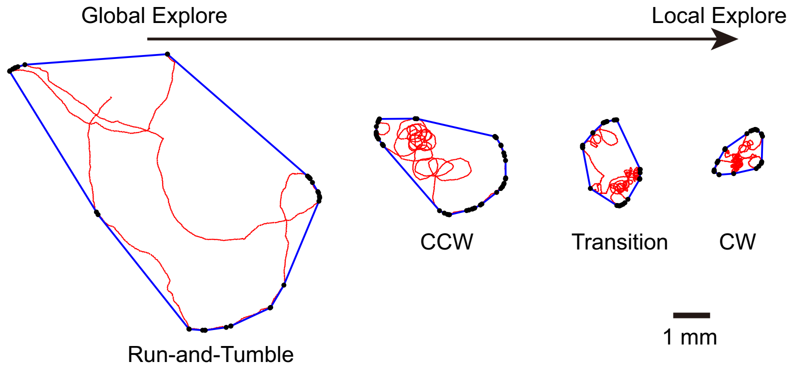


Figure S12: *Chlamydomonas* exhibits various navigation strategy (quantified as the area of 2D convex hull) in different swimming mode. Scale bars, 1 mm.

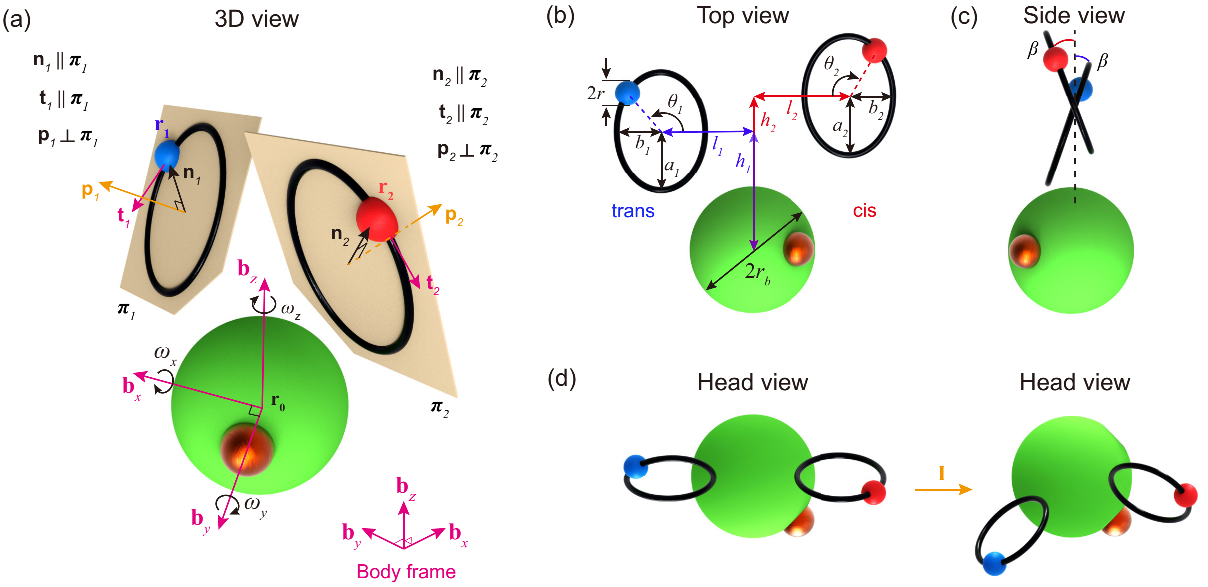


Figure S13: Schematic of the hydrodynamic model of *Chlamydomonas*. (a) 3D view. (b) Top view. (c) Side view. The beat orbits are tilted with an angle $\beta$. (d) Head view.

$$\dot{\mathbf{r}}_i = \dot{\mathbf{r}}_0 + \Omega \times (\mathbf{r}_i - \mathbf{r}_0) + \dot{\mathbf{r}}_i^{'}. \tag{S3}$$

Since the flagellar orbits are elliptical (validated by experiments), $\dot{\mathbf{r}}_i^{'}$ is not aligned with the tangential vector of the orbit $\hat{\mathbf{t}}_i$. According to Kepler's laws, $\dot{\mathbf{r}}_i^{'}$ can be expressed as:

$$\dot{\mathbf{r}}_i^{'} = \frac{dR_i}{dt} \cdot \hat{\mathbf{e}}_{R_i} + R_i \frac{d\theta_i}{dt} \cdot \hat{\mathbf{e}}_{\theta_i} = \dot{R}_i(\theta_i)\dot{\theta}_i \cdot \hat{\mathbf{n}}_i + R_i \dot{\theta}_i \cdot \hat{\mathbf{t}}_i, \tag{S4}$$

where $\hat{\mathbf{n}}_i$ and $\hat{\mathbf{t}}_i$ are unit vectors in normal and tangential direction, respectively. Together with $\hat{\mathbf{p}}_i$, which is perpendicular to the orbit plane $\pi_i$, these vectors form a mutually perpendicular coordinate system describing flagellar beat. The parameter $\dot{\theta}_i$ represents the time rate of change of the phase of flagella beats, while $\dot{R}_i(\theta_i)$ specifies the radial velocity of flagella sphere as a function of the ellipse parameters $a_i$ and $b_i$. Similarly to the experimental waveform (Fig. S5), $a_i$ and $b_i$ are defined to describe the size of the orbit, while $h_i$ and $l_i$ describe the position of the orbit [Fig. S13(b)]. The phase of flagellar beat within a cycle is described by $\theta_i$. Notably, the orbits are tilted by an angle $\beta$, measured to be 0.3 rad [Fig. S13(c)]. Different with previous model [1, 4], here we introduced additional rules to allow the swimmer to dynamically adjust its waveform, driving force, and beat plane angle based on detected light signals [Fig. S13(d)]. For the details, we will discuss in Section 3.4-3.7.

#### 3.2 Hydrodynamic interactions

The flagellar spheres are moved by tangential forces $\mathbf{F}_i^{(t)} = F_i^{(t)}\hat{\mathbf{t}}_i = (1 + c_i \cos(\theta_i + \theta_{i,0}))\hat{\mathbf{t}}_i$. Together with the normal forces $\mathbf{F}_i^{(n)}$ and the constraint forces $\mathbf{F}_i^{(p)}$ (which confine the motion to the orbit plane), the total force on each flagellar sphere is given by $\mathbf{F}_i = F_i^{(t)}(\theta)\hat{\mathbf{t}}_i + F_i^{(n)}\hat{\mathbf{n}}_i + F_i^{(p)}\hat{\mathbf{p}}_i$. The initial flagellar phase is accounted for by $\theta_{i,0}$, which is set as $\theta_{1,0} = \pi$ and $\theta_{2,0} = 7/6\pi$ in all simulations to replicate experimentally observed phase differences.

Hydrodynamic interactions between the spheres representing the cell body and flagella are modeled using the Oseen approximation [9]. The linear relationship between velocities and hydrodynamic forces is expressed as:

$$\dot{\mathbf{r}}_0 = [\mathbf{G}(\mathbf{r}_0 - \mathbf{r}_1) - \gamma_0^{-1}] \cdot \mathbf{F}_1 + [\mathbf{G}(\mathbf{r}_0 - \mathbf{r}_2) - \gamma_0^{-1}] \cdot \mathbf{F}_2, \tag{S5}$$

$$\dot{\mathbf{r}}_1 = \gamma_1^{-1}\mathbf{F}_1 + [\mathbf{G}(\mathbf{r}_1 - \mathbf{r}_2) - \mathbf{G}(\mathbf{r}_1 - \mathbf{r}_0)] \cdot \mathbf{F}_2 - [\mathbf{G}(\mathbf{r}_1 - \mathbf{r}_0)] \cdot \mathbf{F}_1, \tag{S6}$$

$$\dot{\mathbf{r}}_2 = \gamma_2^{-1}\mathbf{F}_2 + [\mathbf{G}(\mathbf{r}_2 - \mathbf{r}_1) - \mathbf{G}(\mathbf{r}_2 - \mathbf{r}_0)] \cdot \mathbf{F}_1 - [\mathbf{G}(\mathbf{r}_2 - \mathbf{r}_0)] \cdot \mathbf{F}_2. \tag{S7}$$

Here, $\gamma_i = 6\pi\eta r$ ($i = 1, 2$), $\gamma_0 = 6\pi\eta r_b$ and $\mathbf{G}(\mathbf{r}) = (\mathbb{I} + \hat{\mathbf{r}}\hat{\mathbf{r}})/8\pi\eta|\mathbf{r}|$, where $\eta$ is the fluid's dynamic viscosity.

At low Reynolds numbers, we impose force-free and torque-free conditions on the swimmer:

$$\mathbf{F}_0 + \mathbf{F}_1 + \mathbf{F}_2 = 0, \tag{S8}$$

$$\mathbf{T}_0 + \mathbf{T}_1 + \mathbf{T}_2 = 0. \tag{S9}$$

Assume that the flagellar sphere radii are much smaller than the body radius ($r \ll r_b$), intrinsic torque from flagellar sphere rotation can be ignored, simplifying the torque-free condition to:

$$8\pi\eta {r_b}^3\Omega + (\mathbf{r}_1 - \mathbf{r}_0) \times \mathbf{F}_1 + (\mathbf{r}_2 - \mathbf{r}_0) \times \mathbf{F}_2 = 0, \tag{S10}$$

where $8\pi\eta {r_b}^3\Omega$ represents the intrinsic torque due to cell body rotation. Equations S3 to S10 form a closed system of equations for describing the 3D dynamics of the cell.

To simplify calculations, we non-dimensionalized the governing equations. Fluid viscosity was normalized by that of water, $\eta \approx 10^{-3}$ pN$\mu m^{-2}$s, while forces were scaled by a typical flagellar beating force, $F \approx 30$ pN [4]. The length scale is the average cell body diameter, $d \approx 10.6$ $\mu$m ($10.6 \pm 0.3$ $\mu$m, $N = 21$ cells).

#### 3.3 Numerical methods

After solving these equations, we calculate the rotation velocities of the spheres at each timestep. Sphere positions are then updated using the quaternion method. For rotation about a unit vector $\hat{\mathbf{y}} = (\omega_x, \omega_y, \omega_z)$ by an angle $\kappa$, the quaternion is:

$$\begin{aligned} q(\hat{\mathbf{y}}, \kappa) &= [q_1, q_2, q_3, q_4] \\ &= [\cos(\kappa/2), \sin(\kappa/2)\omega_x, \sin(\kappa/2)\omega_y, \sin(\kappa/2)\omega_z]. \end{aligned} \tag{S11}$$

The corresponding rotation matrix is:

$$\mathbf{Q} = \begin{bmatrix} 1 - 2{q_3}^2 - 2{q_4}^2 & 2(q_2q_3 - q_1q_4) & 2(q_2q_4 + q_1q_3) \\ 2(q_2q_3 + q_1q_4) & 1 - 2{q_2}^2 - 2{q_4}^2 & 2(q_3q_4 - q_1q_2) \\ 2(q_2q_4 - q_1q_3) & 2(q_3q_4 + q_1q_2) & 1 - 2{q_2}^2 - 2{q_3}^2 \end{bmatrix}. \tag{S12}$$

After rotation, any vector (e.g., $\hat{\mathbf{s}}_t$) is updated as:

$$\hat{\mathbf{s}}_{t+1} = \mathbf{Q}(\hat{\mathbf{y}}, \kappa)\hat{\mathbf{s}}_t. \tag{S13}$$

The swimmer's position is updated iteratively using the fourth-order Runge-Kutta method.

#### 3.4 Orthogonal swimming versus phototaxis

By comparing orthogonal swimming and phototaxis (both with light along the $-z$-axis), we found that orthogonal swimming can be achieved by altering the torque distribution perpendicular to the light plane, which differs from typical phototaxis. For orthogonal swimming [Fig. S14(a)-(d)], the net torques in $x$ and $y$ directions average to zero, while that in $z$ remains nonzero, resulting in a circular trajectory in the $x$–$y$ plane orthogonal to light. In phototaxis [Fig. S14(e)-(h)], nonzero torques in all axes align the cell with the light, preventing orthogonal motion.

In our model, orthogonal swimming results from beat plane reorientation in response to the light signal $S(t)$, realigning net force and torque. Phototaxis results from changing the flagellar driving force ($c_i$) based on light detected by the eyespot [1], described by:

$$S(t) = S_a\,|\mathbf{I}| + S_d(-\mathbf{I} \cdot \hat{\mathbf{e}})\mathrm{H}(-\mathbf{I} \cdot \hat{\mathbf{e}}). \tag{S14}$$

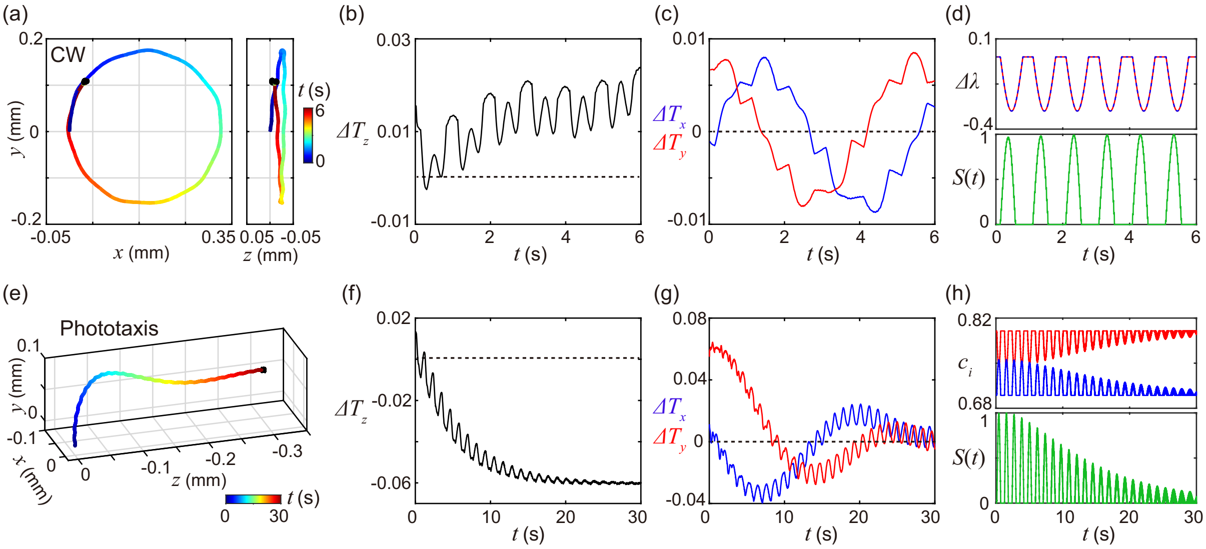


Figure S14: The key of orthogonal swimming is that the integrals of $\Delta T_x$ and $\Delta T_y$ are both zero.

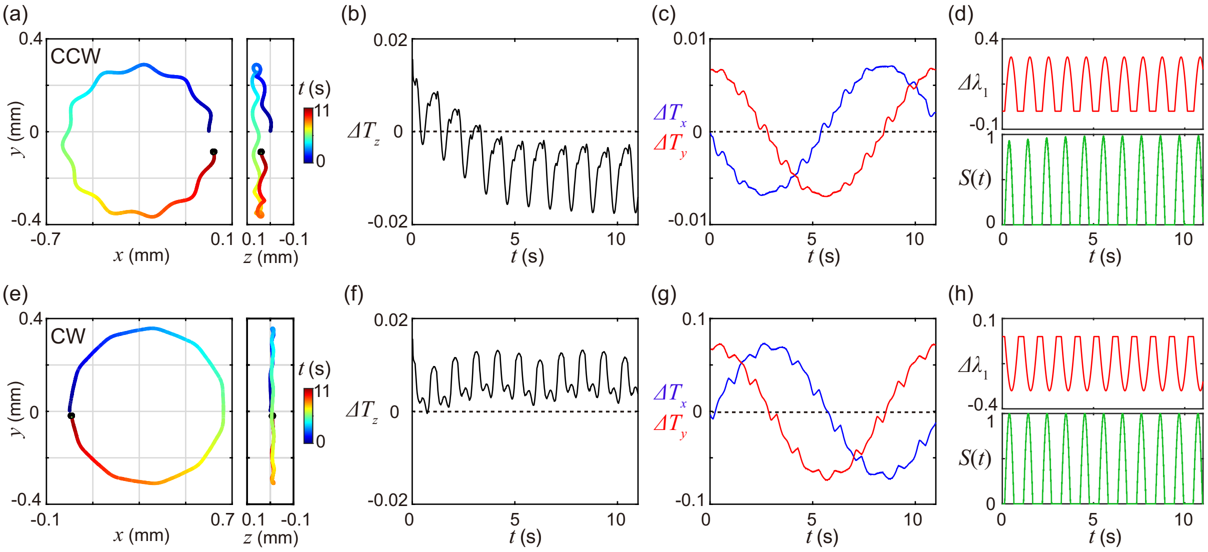


Figure S15: The sign of $\Delta T_z$ determines the handedness of the orthogonal swimming trajectory. $\Delta\lambda_1$ represents the beat plane angle of *trans*-flagellum. Positive $\Delta\lambda$ is defined as reorientation away from the eyespot, and negative $\Delta\lambda$ as toward it.

where $\mathbf{I}$ is the light vector, $\hat{\mathbf{e}}$ is the eyespot direction, $S_a$ and $S_d$ are coupling constants, and H is the Heaviside function. The beat phase is modulated as $c_i = c_0 + K_{c,i}\log(S(t))$. $K_{c,i}$ is a light-dependent coupling constant.

In summary, orthogonal swimming requires net torques orthogonal to light to average to zero, achieved by beat plane reorientation. This difference arises from cell asymmetry: *Chlamydomonas* has two flagella with beat planes oriented ∼45° and ∼135° relative to the eyespot. As the cell rotates, the flagella generate forces not perpendicular to the light, causing the cell to align with light (phototaxis). Our results show that light stimulus can reorient the beat plane to compensate for the intrinsic cell asymmetry, such that the redistributed torque aligns with the orthogonal axis.

#### 3.5 Handedness of orthogonal swimming

Here we examine the factors determining orthogonal swimming handedness. Since the cell moves orthogonally in the $x$–$y$ plane, the net torque along the $z$-axis $\Delta T_z$ determines the trajectory handedness. $\Delta T_z < 0$ produces CCW motion, while $\Delta T_z > 0$ results in CW motion (Fig. S15).

Experimentally, weak light shifts the beat plane away from the eyespot (yielding CCW trajectories), whereas strong light shifts it toward the eyespot (yielding CW trajectories). Simulations reproduce these trends [Fig. S15 (d), (h)]. Notably, handedness switching here is achieved by changing only the beat plane of the *trans*-flagellum, but altering the *cis*-flagellum or both flagella produces the same outcome: away from the eyespot yields CCW, toward the eyespot yields CW.

Thus, the direction of beat plane reorientation under different light intensities determines the sign of $\Delta T_z$ and the handedness of orthogonal swimming. Next, we analyze how the beat plane deflection angle influences the radius of the orthogonal trajectory.

##### 3.6 Radius of orthogonal swimming

To quantify the geometry of orthogonal swimming, we examined how trajectory radius depends on the beat plane reorientation. Simulations reveal an inverse relationship: as the deflection angle increases, the trajectory radius decreases, consistent for both CW and CCW motion.

Using representative cases where the beat plane deflection angle $\Delta\lambda = m \sin(nt)$ (with $m$ from 0.1 to 0.9), we found that larger $m$ values yield smaller radii [Fig. S16(a)-(b)]. Additionally, by varying $m$, the model produces diverse polygonal trajectories [Fig. S16(c)-(h)], with the number of sides roughly proportional to the radius.

These results demonstrate that beat plane dynamics directly control both the size and shape of orthogonal trajectories. Next, we explore whether other subcellular mechanisms, beyond beat plane reorientation, can also modulate the curvature of the orthogonal swimming.

##### 3.7 Curvature modulation by $\Delta h$ and $\Delta\theta$

Previously, we demonstrated that orthogonal swimming is achieved and controlled by beat plane reorientation. We next asked whether other factors can further modulate the curvature of these orthogonal trajectories. To test this, we allow cells to adjust beat extension ($\Delta h$) and beat phase ($\Delta\theta$) by changing flagellar driving force ($c_i$) in response to detected light. The flagella beat extensions and beat phases are modulated by: $h_i = h_0 + K_{h,i} \log(S(t))$ and $c_i = c_0 + K_{c,i} \log(S(t))$ in CCW and CW modes respectively ($i = 1, 2$ for *trans*- and *cis*-flagellum). $K_{h,i}$ and $K_{c,i}$ are light-dependent coupling constants.

Simulations reveal that modulating beat phase or extension, in addition to beat plane orientation, enables further control of trajectory curvature. When only the beat plane is adjusted, trajectories are smooth and circular, but do not match the curvature change rate seen experimentally [Fig. S17(a)-(c) and Fig. S18(a)-(c)]. Introducing differences in flagellar driving force ($c_i$; Fig. S17) or beat extension ($h_i$; Fig. S18) reproduces more complex patterns, such as flower-like trajectories and especially matches the dynamic curvature changes.

Thus, while beat extension and phase cannot generate orthogonal swimming on their own, they may provide additional flexibility in tuning trajectory curvature once the beat plane is reoriented. This dynamic interplay allows the cell to finely adjust its swimming path in response to light, reflecting the adaptability of its sensor–actuator system. In the next section, we summarize our main findings and discuss their implications for understanding light-guided swimming in *Chlamydomonas*.

##### 3.8 Summary

Using detailed hydrodynamic simulations, we systematically investigated how *Chlamydomonas* cells achieve and regulate orthogonal swimming under light.

(1) Torque distribution: Orthogonal swimming arises when net torques orthogonal to the light direction integrate to zero—achieved by reorienting the flagellar beat plane.

(2) Handedness control: The sign of the residual torque along the light direction ($\Delta T_z$) determines whether the trajectory is CW or CCW. This is set by the direction of beat plane reorientation: low light induces CCW motion (beat plane away from eyespot), while high light induces CW motion (towards eyespot), matching experiments.

(3) Geometric properties: The radius of the orthogonal trajectory is inversely related to the amplitude of beat plane deflection. By tuning this, cells generate a range of trajectory shapes, including polygons

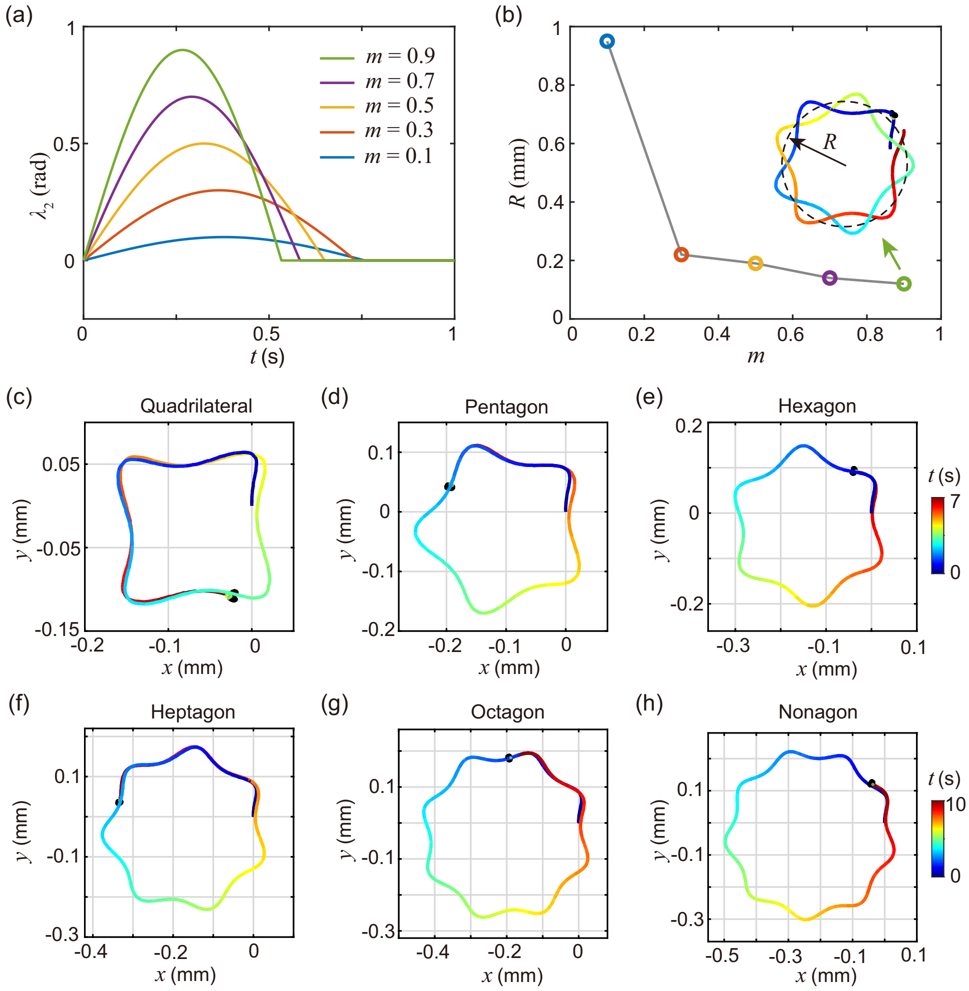


Figure S16: By increasing the beat plane reorientation angle, both the radius and the shape of the cell's trajectory are modulated. (a) Time course of the beat plane reorientation angle ($\Delta\lambda$). (b) Trajectory radius for different $m$ values. (c–g) Different $m$ values generate polygonal trajectories with varying numbers of sides. The swimming duration is 7 seconds for (c–e) and 10 seconds for (f–h).

with different numbers of sides.

(4) Multiple pathways for curvature modulation: Beat plane reorientation is essential for orthogonal swimming; subsequently, light-induced changes in beat phase ($\Delta\theta$) and beat extension ($\Delta h$) can further modulate the curvature of the orthogonal circular trajectory, enabling flexible adaptation of swimming patterns to environmental cues.

Taken together, these results provide a comprehensive mechanistic understanding of light-guided swimming in *Chlamydomonas*. Our model reproduces key experimental observations (handedness switching, trajectory types) and predicts how other subcellular changes enable adaptive navigation.

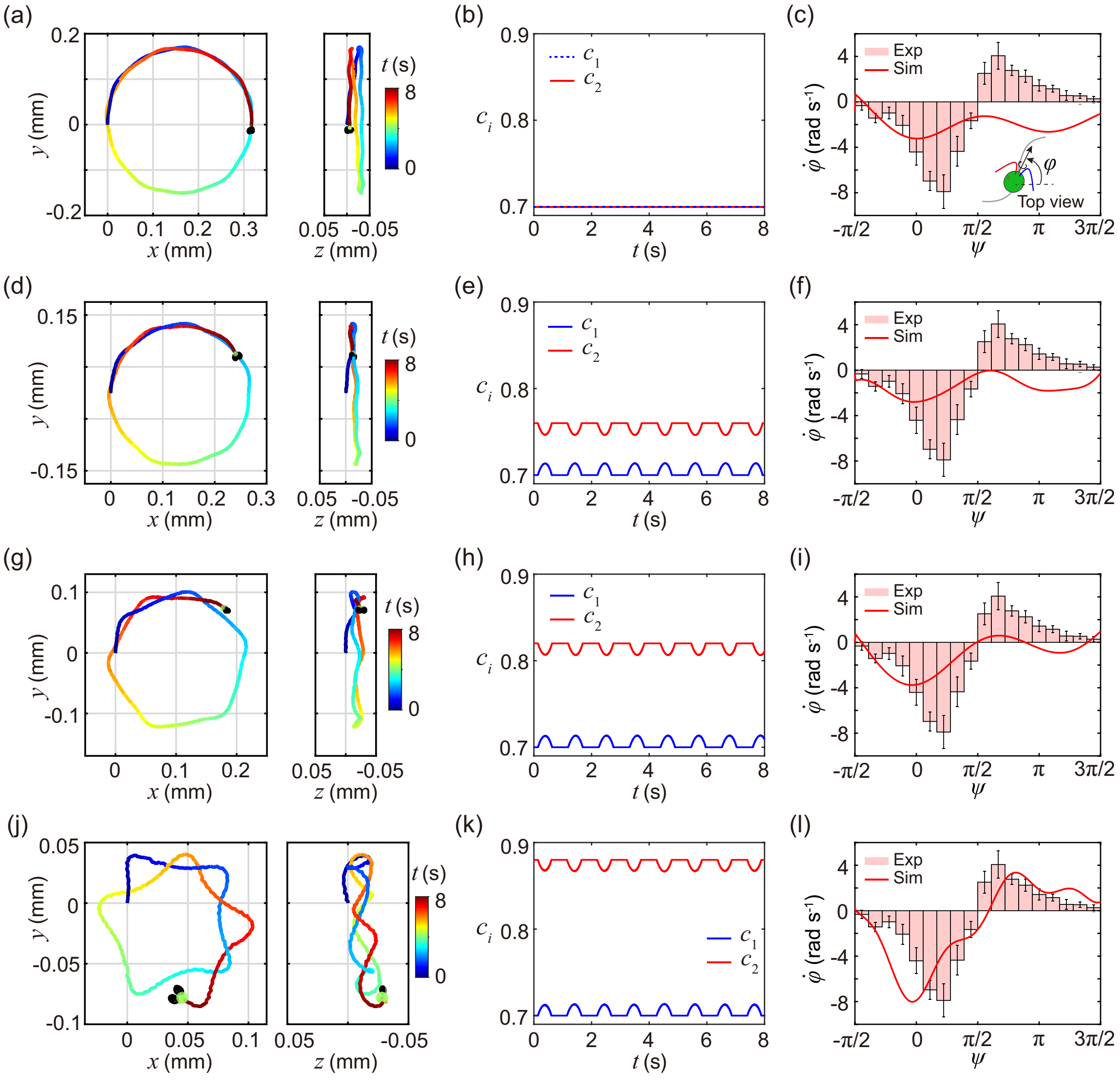


Figure S17: $\Delta\theta$ can contribute to curvature modulation in *Chlamydomonas*. The flagellar waveform and beat plane dynamics followed experimental measurements, whereas $c_1$ and $c_2$ were further adjusted in response to the light signal in (d)–(l). Initial value without light modulation: (a–c) $c_1 = c_2 = 0.7$; (d–f) $c_1 = 0.7$, $c_2 = 0.76$; (g–i) $c_1 = 0.7$, $c_2 = 0.82$; and (j–l) $c_1 = 0.7$, $c_2 = 0.88$.

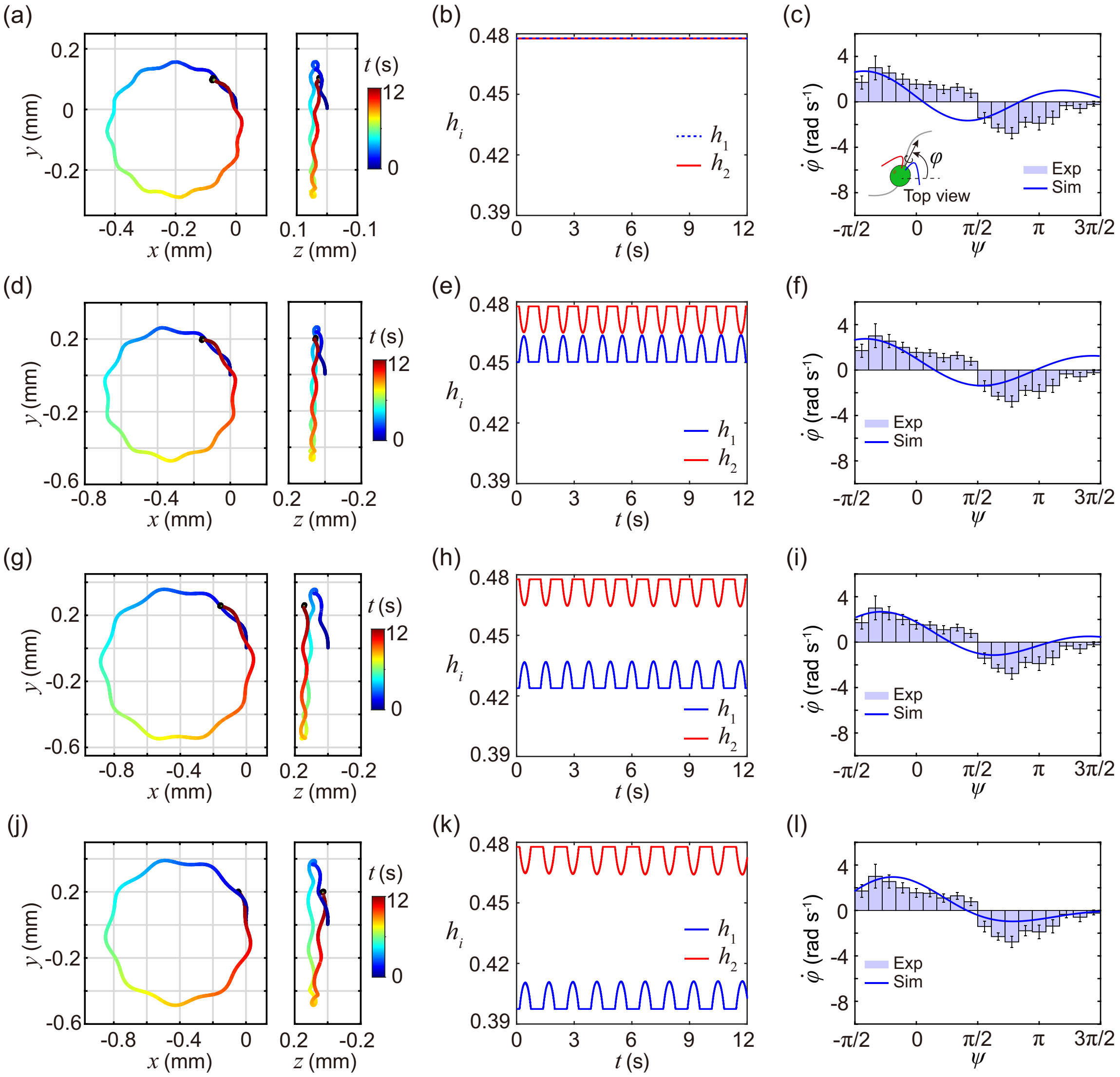


Figure S18: $\Delta h$ can contribute to curvature modulation in *Chlamydomonas*. The flagellar waveform and beat plane dynamics followed experimental measurements, whereas $h_1$ and $h_2$ were further adjusted in response to the light signal in (d)–(l). Initial value without light modulation: (a–c) $h_1 = h_2 \approx 0.479$; (d–f) $h_1 \approx 0.479$, $h_2 \approx 0.466$; (g–i) $h_1 \approx 0.479$, $h_2 \approx 0.453$; (j–l) $h_1 \approx 0.479$, $h_2 \approx 0.40$.
